# sangerFlow, a Sanger sequencing-based bioinformatics pipeline for pests and pathogens identification

**DOI:** 10.1101/2024.05.10.593518

**Authors:** M. Asaduzzaman Prodhan, Matthew Power, Monica Kehoe

## Abstract

Sequencing of a Polymerase Chain Reaction product (amplicon) is called amplicon sequencing. Amplicon sequencing allows for reliable identification of an organism by amplifying, sequencing, and analysing a single conserved marker gene or DNA barcode. As this approach generally involves a single gene, it is a light-weight protocol compared to multi-locus or whole genome sequencing for diagnostic purposes; yet considerably reliable. Therefore, Sanger-based high-quality amplicon sequencing is widely deployed for species identification and high-throughput biosecurity surveillance. However, keeping up with the data analysis in a large-scale surveillance or diagnostic settings could be a limiting factor because it involves manual quality control of the raw sequencing data, alignment of the forward and reverse reads, and finally web-based Blastn search of all the amplicons. Here, we present a bioinformatics pipeline that automates the entire analysis. As a result, the pipeline is scalable with high-volume of samples and reproducible. Furthermore, the pipeline leverages the modern open-source Nextflow and Singularity concept, thus it does not require software installation except Nextflow and Singularity, software subscription, or programming expertise from the end users making it widely adaptable.

**Availability and implementation:** sangerFlow source code and documentation are freely available for download at GitHub, implemented in Nextflow and Singularity.

## Introduction

Sanger sequencing^1^ of single-target PCR product (amplicon) is widely used as a low-cost molecular diagnostic tool^2^. For diagnostic purposes, it is primarily used to identify a species or microorganism by performing a sequence similarity analysis using National Center for Biotechnology Information (NCBI)’s Basic Local Alignment Search Tool for nucleotide (Blastn)^3^. For example, mitochondrial CO1 for insect species identification^4^, 16S rRNA for pathogenic bacterial identification or 28S Internal Transcribed ribosomal gene Spacer (28S–ITS) for fungal identification^5^. The Sanger sequencing data of the forward and reverse reads can be analysed using Graphical User Interface (GUI)-based commercial software such as Geneious^6^ and CLCbio^7^ or non-commercial software like SnackVar^8^, GLASS^9^, and SeqTrace^10^. While these GUI-based tools are generally user-friendly, they are designed for manual processing of the data. In a large-scale biosecurity surveillance setting, the GUI-based data processing could be a time- and resource-limiting factor. However, some command line based open-source tools such SangeR^11^, ASAP^12^ have been developed to analyse sanger sequencing data. However, SangeR^11^ has been developed to perform mutation analysis while ASAP^12^ can perform protein alignment. To the best of our knowledge, command line based open-source tools that automatically processes Sanger sequencing raw data and performs a Blastn search to identify the subject organism are rare. Although such a tool is in high demand for a biosecurity incident where wide-ranging surveillance is underway so that i) large volumes of samples can be processed rapidly, and ii) species identification can be streamlined. Here, we present a bioinformatics pipeline based on Nextflow and Singularity called sangerFlow. sangerFlow takes Sanger amplicon sequencing raw data and returns the targeted results i.e., the Blastn hits for each sample. Furthermore, the Blastn hits are sorted based on the highest bitscore and lowest e-value and produced in a concise Tab-Separated Values (TSV) format per sample for the convenience of quick species identification. In addition, the pipeline leverages Nextflow and Singularity containers. Therefore, the benefits of sangerFlow also include no-installation of software except for Nextflow and Singularity. Furthermore, the pipeline scripts are hosted in GitHub (https://github.com/asadprodhan/sangerFlow) keeping them away from the general users. This makes sangerFlow less dependent on users’ programming expertise so that it can be implemented easily with minimal personnel training. All these features make the pipeline ideal for large-scale Sanger sequencing data analysis and user-friendly.

## Materials and Methods

### Workflow description

Figure 1 shows the sangerFlow workflow. The pipeline takes both .seq and .fasta file from Sanger sequencing as input data. The forward and reverse reads are fed into the workflow as two individual data channels. In the forward-read channel, the reads are re-named as per their sample names using awk commands followed by trimming of the ambiguous nucleotides using sed commands. In the reverse-read channel, the reverse reads are re-named as per their samples, converted to their reverse-complement strand using bioawk^13^, and trimmed for ambiguous nucleotides. Then, the trimmed reads are concatenated before subjecting them to the pairwise alignment using Clustal Omega^14^. The most representative sequence is extracted as .fasta file from the alignment file using The European Molecular Biology Open Software Suite (EMBOSS) Cons^15^. The fasta-headers are re-named with sample names before passing them as query sequence to the Blastn search^16^. The pipeline produces Blastn hits as xml, html and tsv file formats for the convenience of visualising the Blastn results. xml files are compatible to visualise in MEGAN^17^ while the html files can be opened in any internet browser. The tsv files present the Blastn results for individual samples in the order of highest to lowest similarities. In addition, the pipeline compiles a master Blastn result sheet comprising the user-defined topmost Blastn hits of all the samples that allows for quick checking of the results. Users can also set the e-value and cpu numbers for the Blastn analysis as described in the following section as well as in https://github.com/asadprodhan/sangerFlow.

**Figure 1.**
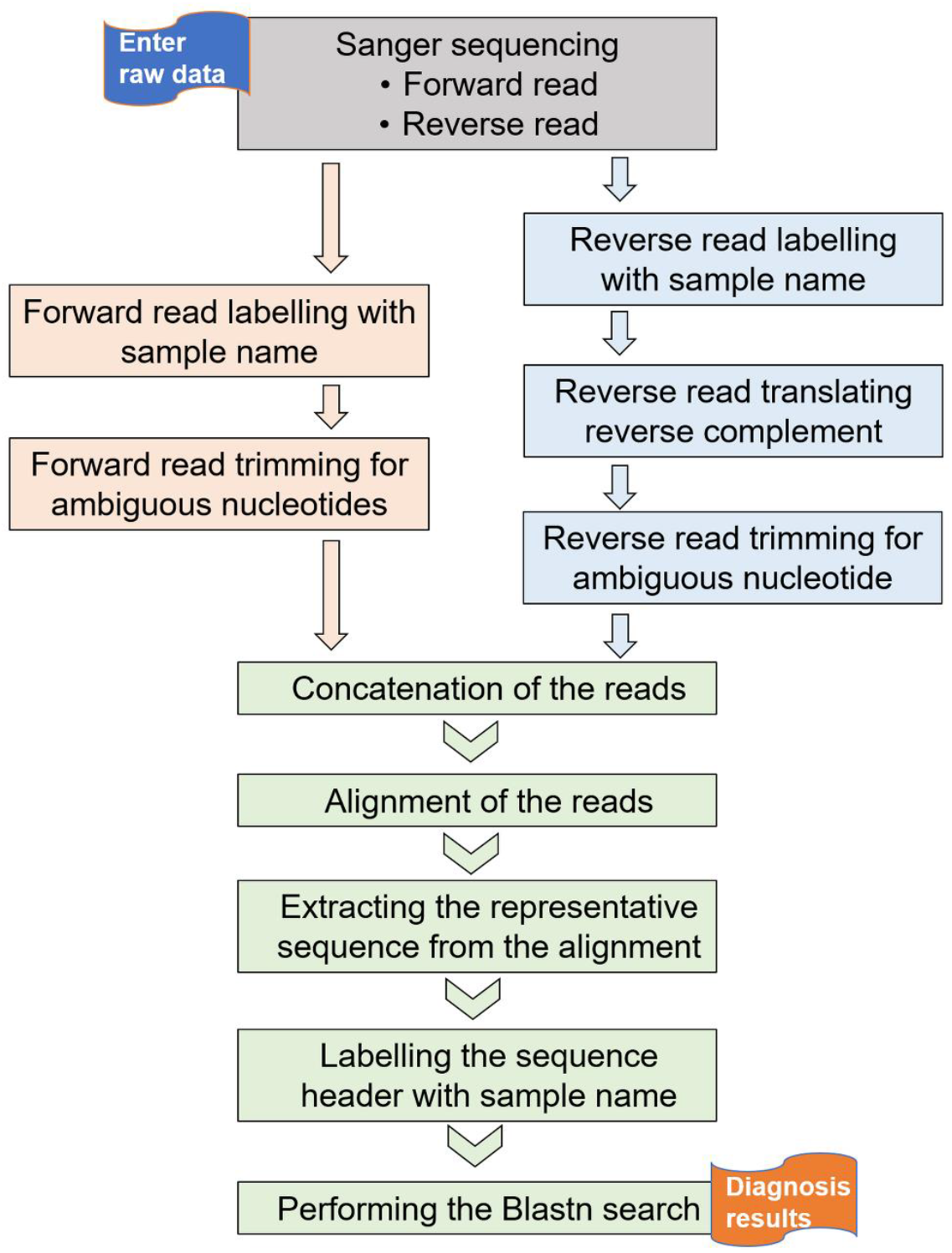
sangerFlow.

### Software implementation and availability

sangerFlow is written using the Nextflow workflow manager coupled with Singularity containers. Therefore, sangerFlow can be run with only two softwares-Nextflow and Singularity-installed on the working computer. The pipeline requires only one script called main.nf and one resources configuration file called nextflow.config. The main.nf script contains all the codes to carry out the individual analyses in a step-wise fashion. The nextflow.config contains Singularity container links for each computational analysis. Furthermore, to simplify the implementation of the workflow, main.nf and nextflow.config are hosted in GitHub (https://github.com/asadprodhan/sangerFlow). Consequently, the sangerFlow can be implemented with only the following coding snippet: nextflow run asadprodhan/sangerFlow -r VERSION-NUMBER --evalue=User_defined --topHits= User_defined --cpus= User_defined --db=“/path/to/your/blastn_database”

The sangerFlow workflow is accompanied with an intuitive user guide hosted in sangerFlow GitHub page (https://github.com/asadprodhan/sangerFlow). The results directory contains the final results while the temp directory includes all the intermediate files generated by sangerFlow in case any sample requires manual inspection for clarity.

### Use case

We have successfully applied sangerFlow to process hundreds of surveillance samples in-house. To demonstrate the utility here, we ran sangerFlow with two publicly available datasets. The test datasets can be found in Table 1 and 2. To run sangerFlow, the users just supply the data and run sangerFlow (nextflow run asadprodhan/sangerFlow -r 18e5299 --evalue=0.05 --topHits=1 --cpus=16 -- db=“/media/Disk/blastn_db”) as depicted in Fig. 2 (Step 1 & 2, respectively). sangerFlow will be automatically launched from its GitHub repository and process all the data supplied without any manual intervention. In the end, there will be three new directories called ‘results’, ‘temp, and ‘work’ (Fig. 2, Step 3). All the Blastn hits will be in the ‘results’ directory. Temp directory will contain the intermediate files like contigs in case some samples require revisiting the intermediate files. The ‘work’ directory is auto-generated by Nextflow that can be discarded.

**Table 1.**
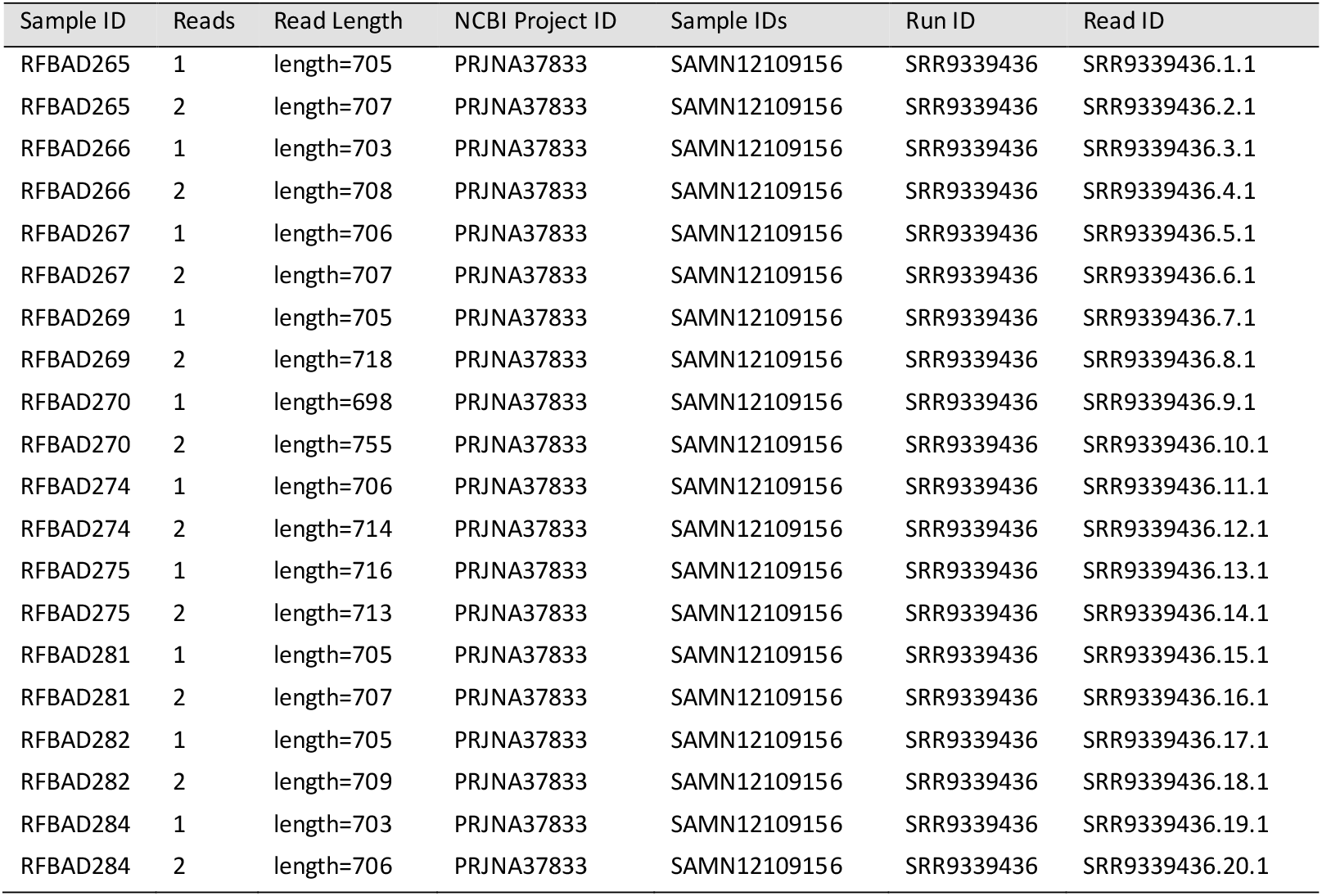
Reference of test dataset 1.

| Sample ID | Reads | Read Length | NCBI Project ID | Sample IDs | Run ID | Read ID |
| --- | --- | --- | --- | --- | --- | --- |
| RFBAD265 | 1 | length=705 | PRJNA37833 | SAMN12109156 | SRR9339436 | SRR9339436.1.1 |
| RFBAD265 | 2 | length=707 | PRJNA37833 | SAMN12109156 | SRR9339436 | SRR9339436.2.1 |
| RFBAD266 | 1 | length=703 | PRJNA37833 | SAMN12109156 | SRR9339436 | SRR9339436.3.1 |
| RFBAD266 | 2 | length=708 | PRJNA37833 | SAMN12109156 | SRR9339436 | SRR9339436.4.1 |
| RFBAD267 | 1 | length=706 | PRJNA37833 | SAMN12109156 | SRR9339436 | SRR9339436.5.1 |
| RFBAD267 | 2 | length=707 | PRJNA37833 | SAMN12109156 | SRR9339436 | SRR9339436.6.1 |
| RFBAD269 | 1 | length=705 | PRJNA37833 | SAMN12109156 | SRR9339436 | SRR9339436.7.1 |
| RFBAD269 | 2 | length=718 | PRJNA37833 | SAMN12109156 | SRR9339436 | SRR9339436.8.1 |
| RFBAD270 | 1 | length=698 | PRJNA37833 | SAMN12109156 | SRR9339436 | SRR9339436.9.1 |
| RFBAD270 | 2 | length=755 | PRJNA37833 | SAMN12109156 | SRR9339436 | SRR9339436.10.1 |
| RFBAD274 | 1 | length=706 | PRJNA37833 | SAMN12109156 | SRR9339436 | SRR9339436.11.1 |
| RFBAD274 | 2 | length=714 | PRJNA37833 | SAMN12109156 | SRR9339436 | SRR9339436.12.1 |
| RFBAD275 | 1 | length=716 | PRJNA37833 | SAMN12109156 | SRR9339436 | SRR9339436.13.1 |
| RFBAD275 | 2 | length=713 | PRJNA37833 | SAMN12109156 | SRR9339436 | SRR9339436.14.1 |
| RFBAD281 | 1 | length=705 | PRJNA37833 | SAMN12109156 | SRR9339436 | SRR9339436.15.1 |
| RFBAD281 | 2 | length=707 | PRJNA37833 | SAMN12109156 | SRR9339436 | SRR9339436.16.1 |
| RFBAD282 | 1 | length=705 | PRJNA37833 | SAMN12109156 | SRR9339436 | SRR9339436.17.1 |
| RFBAD282 | 2 | length=709 | PRJNA37833 | SAMN12109156 | SRR9339436 | SRR9339436.18.1 |
| RFBAD284 | 1 | length=703 | PRJNA37833 | SAMN12109156 | SRR9339436 | SRR9339436.19.1 |
| RFBAD284 | 2 | length=706 | PRJNA37833 | SAMN12109156 | SRR9339436 | SRR9339436.20.1 |

**Table 2.** Reference of test dataset 2 (chromatograms)

| Name (Reference Link) | Sample ID (BOLD) | BOLD BIN (Cluster ID) | Taxonomic Identity | Collection Location |
| --- | --- | --- | --- | --- |
| LHMD002-07<br>https://v3.boldsystems.org/index.php/Public_RecordView?processid=LHMD002-07 | RNP-06-0451-02.2 | BOLD:AAB9578 | *Xyleborinus saxesenii* | British Columbia, Canada |
| LHMD004-07<br>https://v3.boldsystems.org/index.php/Public_RecordView?processid=LHMD004-07 | RNP-06-0451-02.4 | BOLD:AAB9577 | *Xyleborinus saxesenii* | British Columbia, Canada |
| LHMD014-07<br>https://v3.boldsystems.org/index.php/Public_RecordView?processid=LHMD014-07 | RNP-06-0001-03.4 | BOLD:AAB9577 | *Xyleborinus saxesenii* | British Columbia, Canada |

**Table 3:**
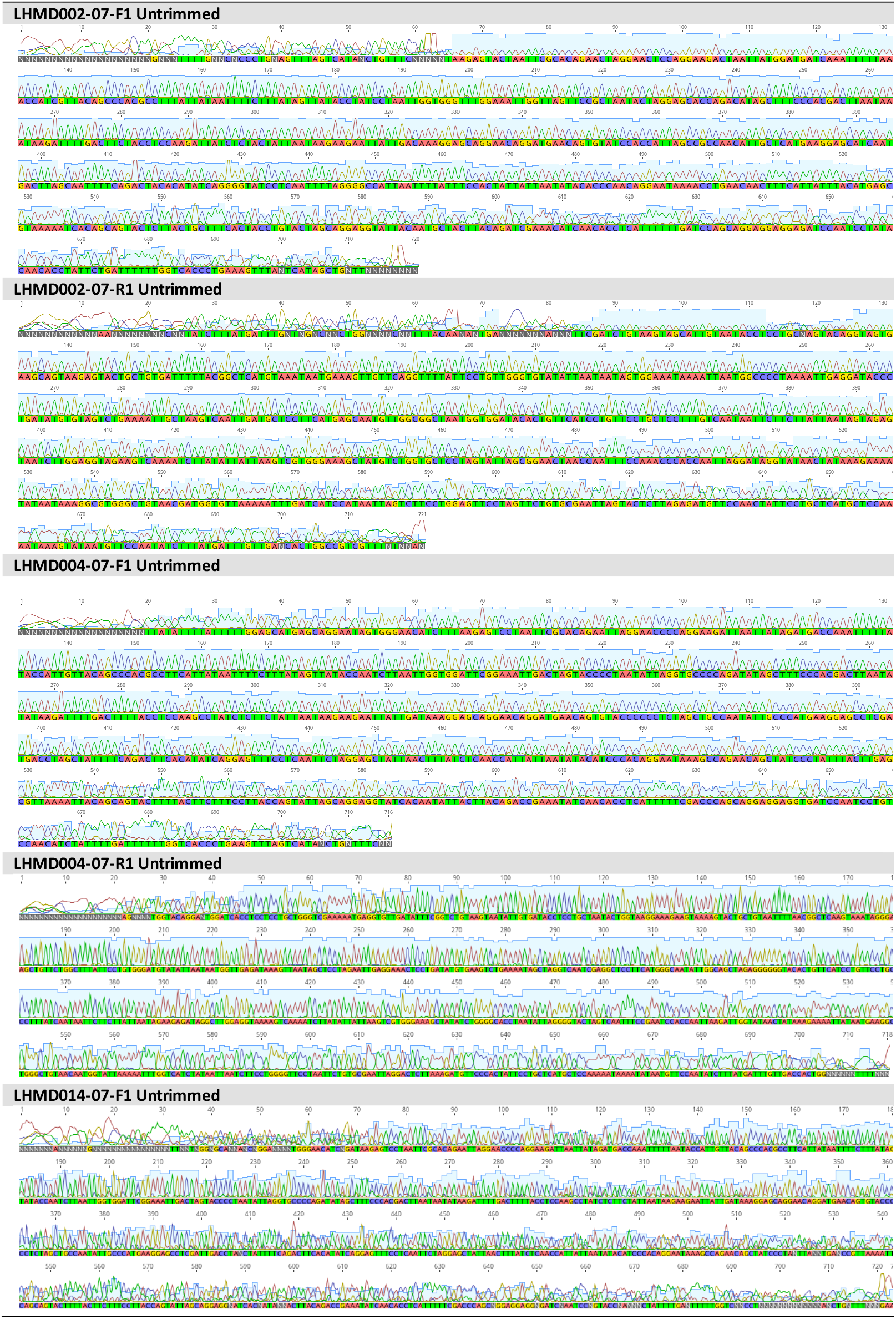
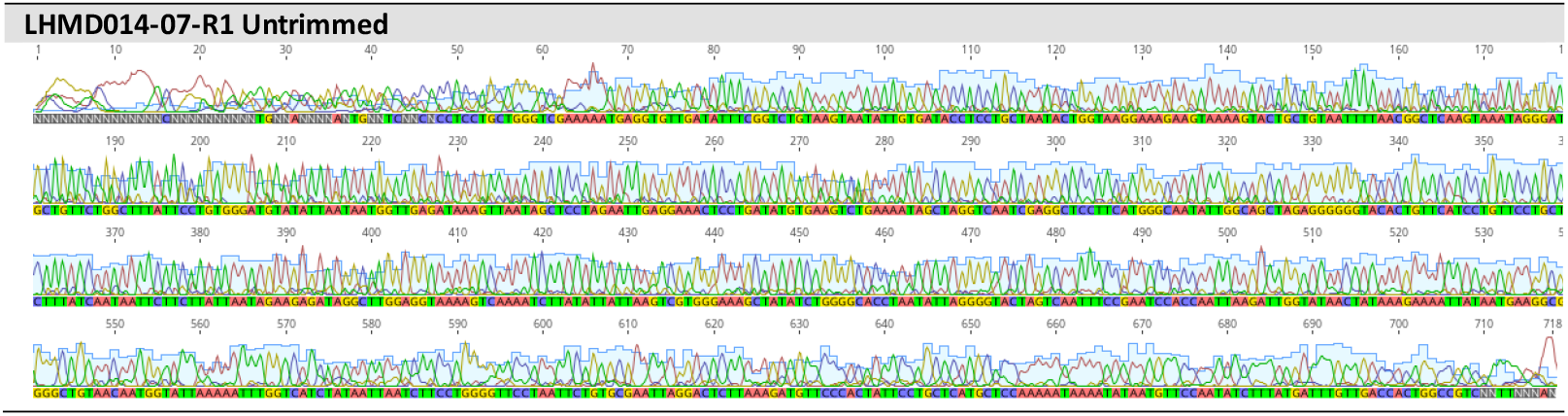
Sanger sequencing chromatograms for test dataset 2.

**Table 4.**
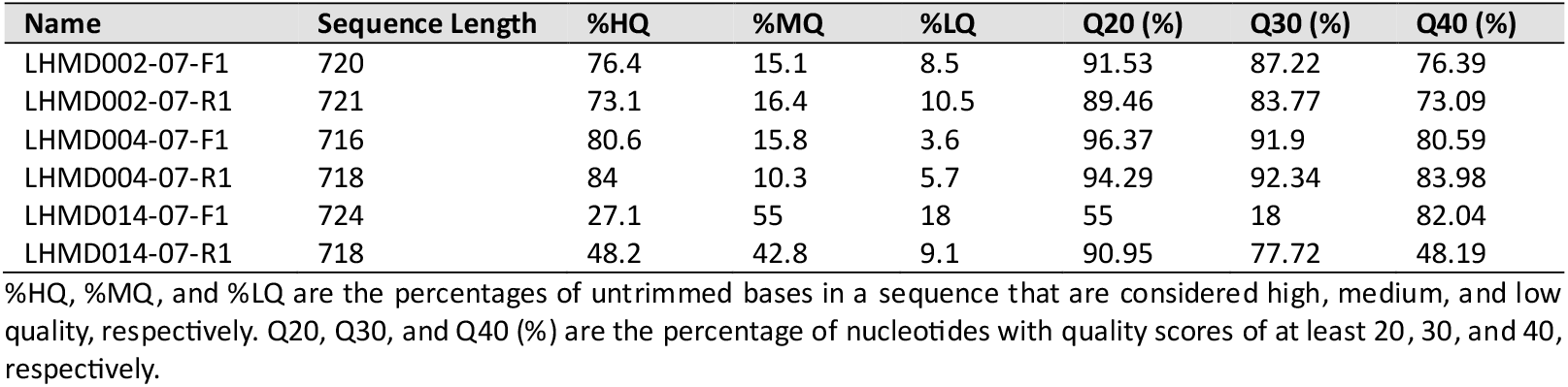
Sequencing quality stats of the reference dataset 2 (chromatograms)

| Name | Sequence Length | %HQ | %MQ | %LQ | Q20 (%) | Q30 (%) | Q40 (%) |
| --- | --- | --- | --- | --- | --- | --- | --- |
| LHMD002-07-F1 | 720 | 76.4 | 15.1 | 8.5 | 91.53 | 87.22 | 76.39 |
| LHMD002-07-R1 | 721 | 73.1 | 16.4 | 10.5 | 89.46 | 83.77 | 73.09 |
| LHMD004-07-F1 | 716 | 80.6 | 15.8 | 3.6 | 96.37 | 91.9 | 80.59 |
| LHMD004-07-R1 | 718 | 84 | 10.3 | 5.7 | 94.29 | 92.34 | 83.98 |
| LHMD014-07-F1 | 724 | 27.1 | 55 | 18 | 55 | 18 | 82.04 |
| LHMD014-07-R1 | 718 | 48.2 | 42.8 | 9.1 | 90.95 | 77.72 | 48.19 |

%HQ, %MQ, and %LQ are the percentages of untrimmed bases in a sequence that are considered high, medium, and low quality, respectively. Q20, Q30, and Q40 (%) are the percentage of nucleotides with quality scores of at least 20, 30, and 40, respectively.

**Figure 2:**
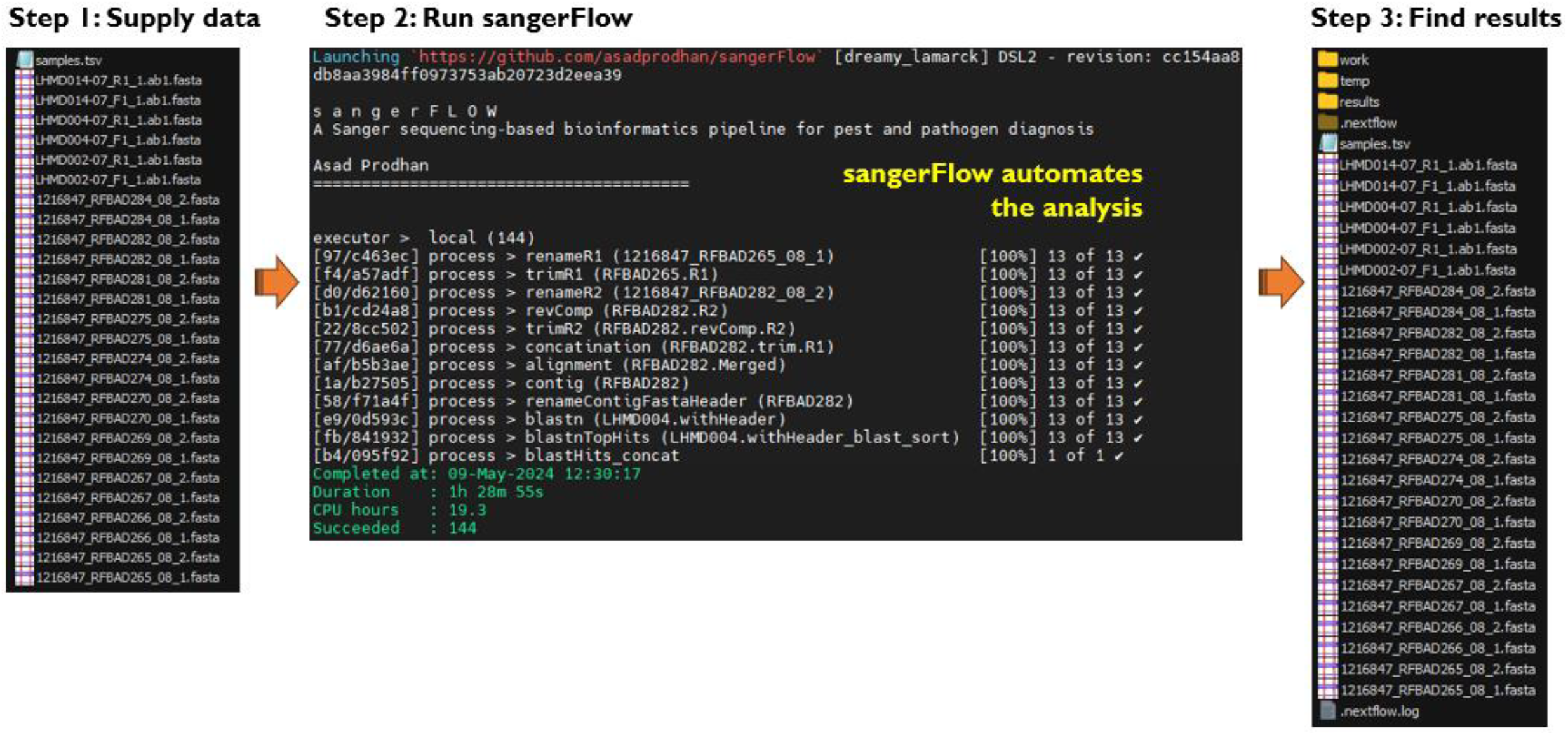
Execution of sangerFlow. For details, see the user protocol at https://github.com/asadprodhan/sangerFlow

## Results

SangerFlow is a command line-based bioinformatics tool that is designed to automatically analyse the Sanger sequencing data of PCR amplicons. Our state-wide biosecurity surveillance for exotic pests, often uses PCR assays resulting in a large influx of Sanger PCR amplicon sequencing data per day. This warrants a high-throughput analysis and led to developing this pipeline, sangerFlow. sangerFlow automatically carried out data quality control, analysis, and Blastn search. Consequently, our species identification was streamlined. We validated the accuracy of the pipeline by comparing the results from the pipeline and those from manual analysis.

To demonstrate the sangerFlow performance here, we used two test datasets that were comprised of PCR Sanger sequencing forward and reverse reads. First, we manually removed the ambiguous nucleotides from the forward and reverse sequences using Geneious^6^ (Table 5), aligned them, extracted the consensus sequence, and finally searched them in the NCBI database using the web Blastn^16^ via Geneious^6^. Then, we ran sangerFlow pipeline on the fasta files of the same datasets that automatically provided with Blastn outputs for each sample (Table 6). However, as the input and output files in sangerFlow were fasta format, there were no visualisation of the trimmed sequences. Finally, we compared the sangerFlow-derived Blastn outputs with those derived from manual processing (Table 7). The comparison of the results demonstrated consistency between manual analysis and sangerFlow (Table 7).

**Table 5:**
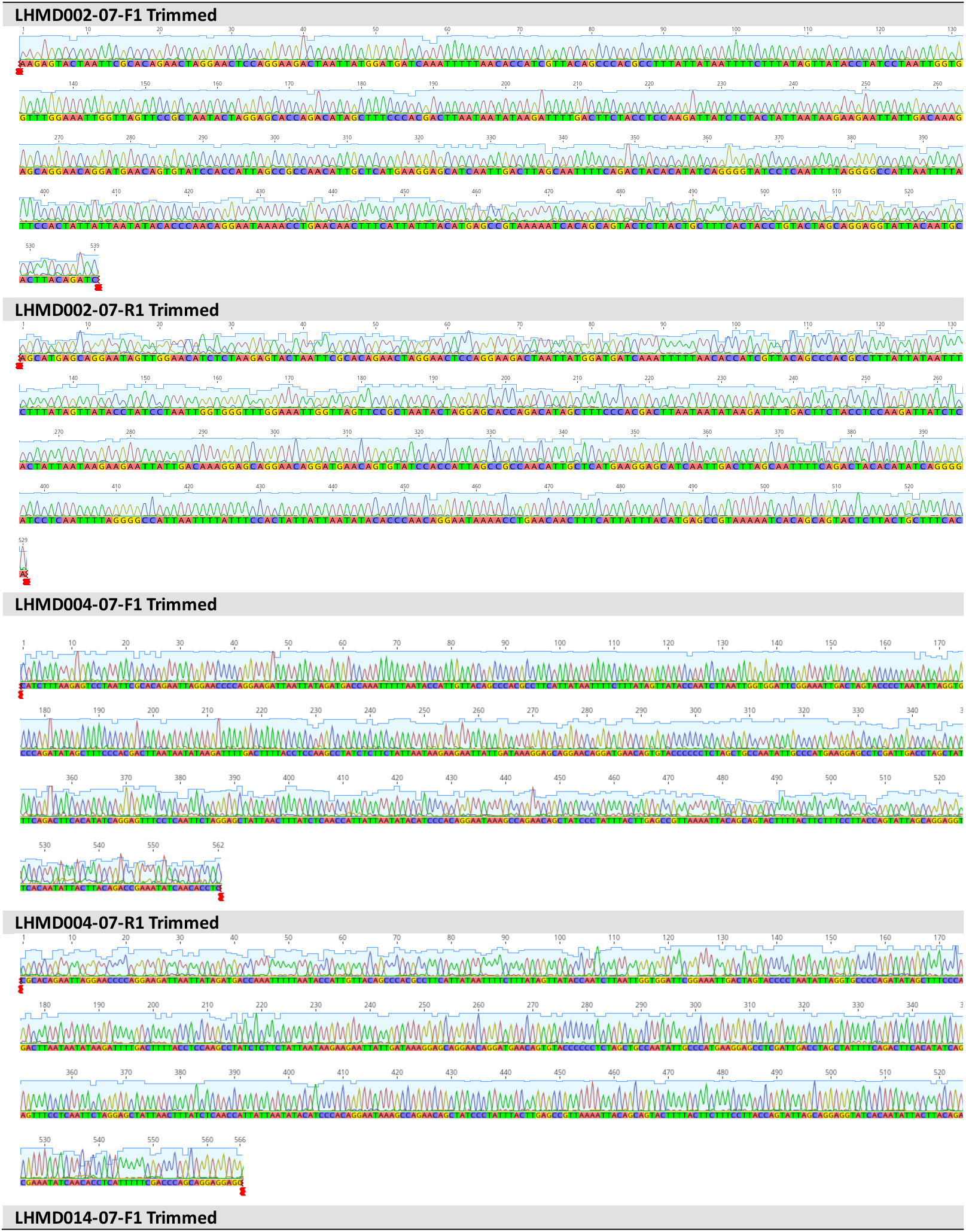
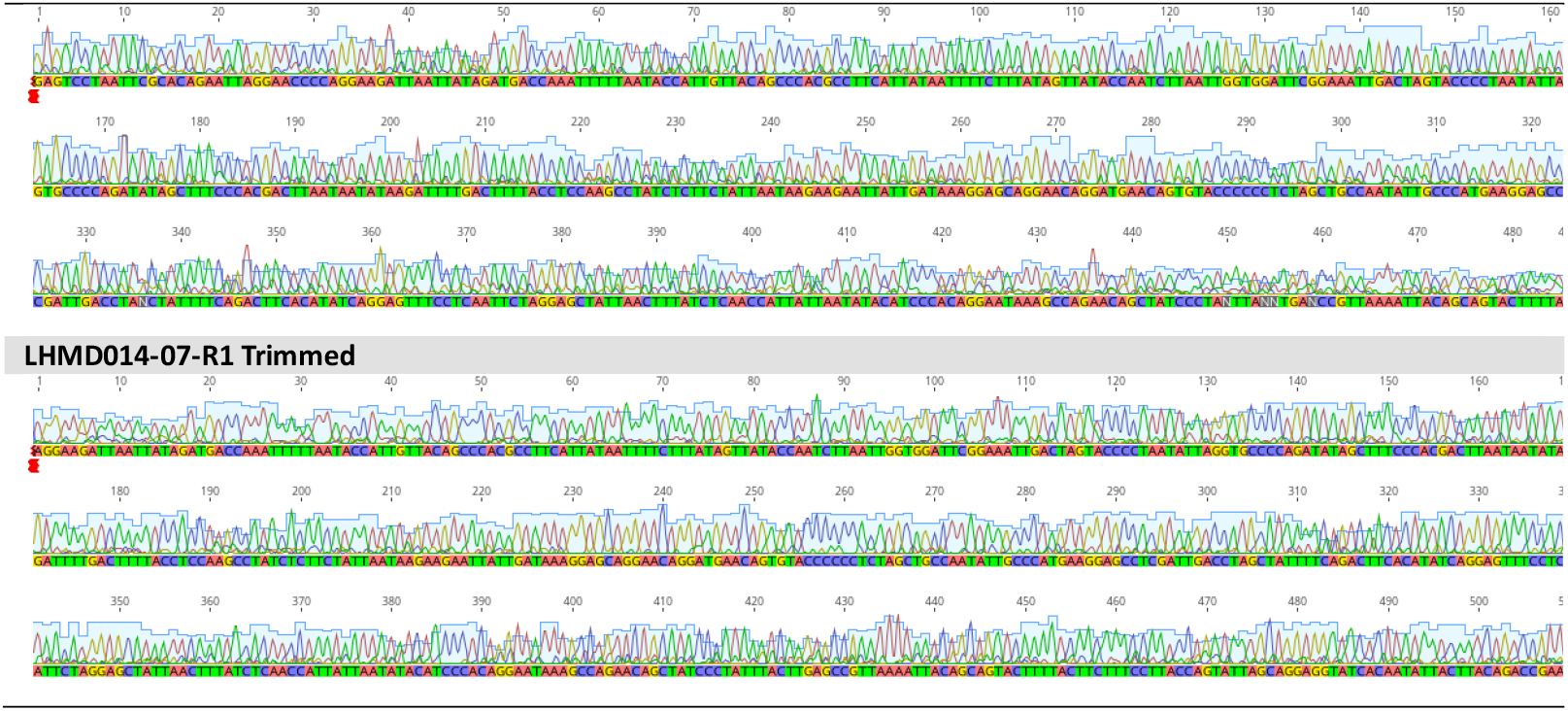
Manually quality-controlled Sanger sequencing chromatograms for test dataset 2.

**Table 6:**
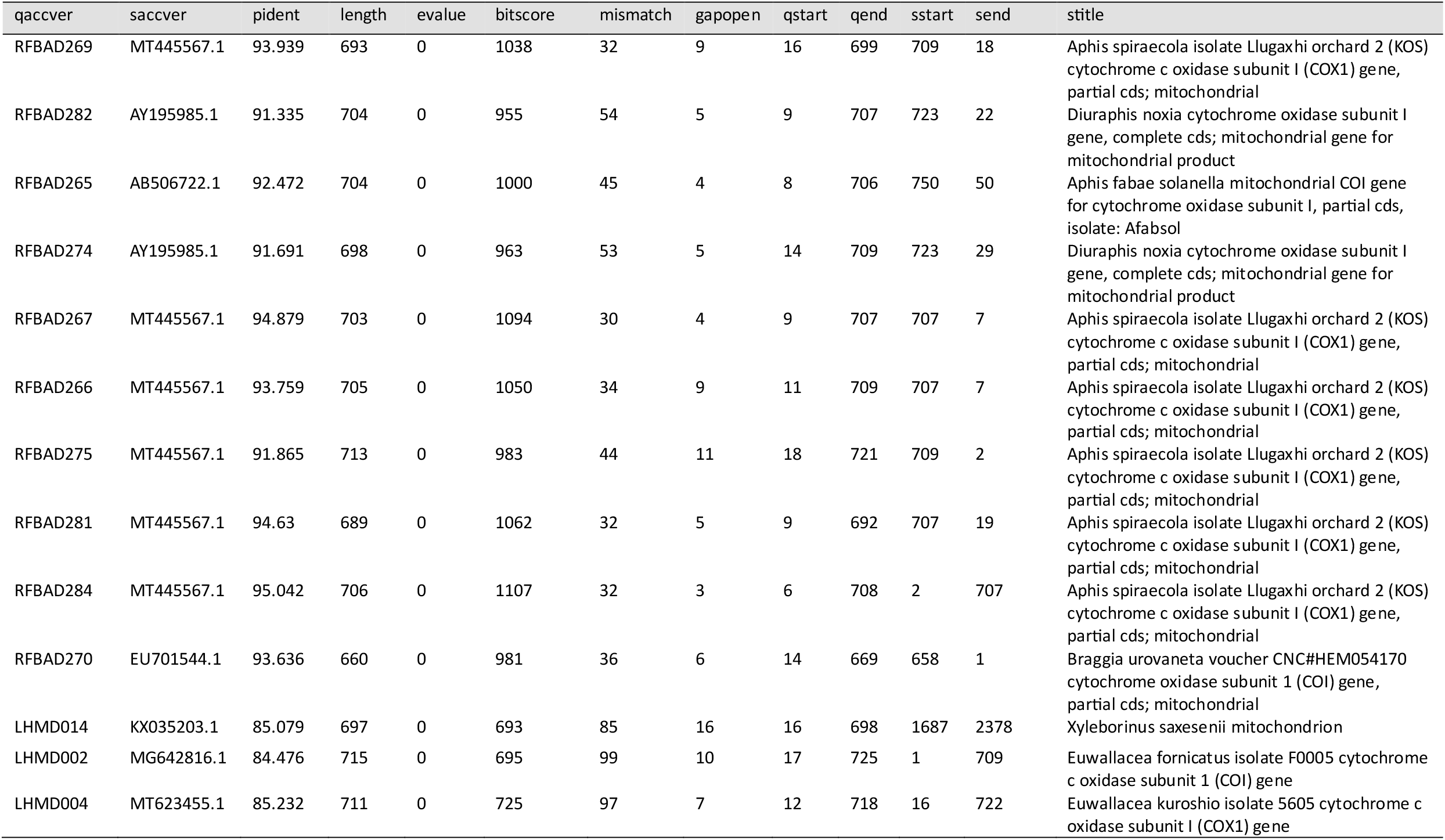
sangerFlow result sheet containing the Blastn IDs of all samples in the run.

**Table 7.** Comparison of taxa Ids derived from sangerFlow and manual analysis.

| sangerFlow |  |  | Geneious |  |
| --- | --- | --- | --- | --- |
| Sample ID | Pairwise % | Taxa ID | Pairwise % | Taxa ID |
| RFBAD265 | 99.50% | *Aphis pomi* | 100% | *Aphis pomi* |
| RFBAD266 | 97.90% | *Aphis pomi* | 100% | *Aphis pomi* |
| RFBAD267 | 98.60% | *Aphis pomi* | 99.90% | *Aphis pomi* |
| RFBAD269 | 99.50% | *Aphis pomi* | 100% | *Aphis pomi* |
| RFBAD270 | 99.70% | *Aphis pomi* | 100% | *Aphis pomi* |
| RFBAD274 | 99.70% | *Aphis pomi* | 100% | *Aphis pomi* |
| RFBAD275 | 96.40% | *Aphis pomi* &<br>*Aphis spiraecola* | 99.80% | *Aphis pomi* &<br>*Aphis spiraecola* |
| RFBAD281 | 99.50% | *Aphis pomi* | 100% | *Aphis pomi* |
| RFBAD282 | 99.20% | *Aphis pomi* | 100% | *Aphis pomi* |
| RFBAD284 | 99.50% | *Aphis pomi* | 99.90% | *Aphis pomi* |
| LHMD002-07 | 100 | *Xyleborinus saxesenii* | 100 | *Xyleborinus saxesenii* |
| LHMD004-07 | 100 | *Xyleborinus saxesenii* | 100 | *Xyleborinus saxesenii* |
| LHMD014-07 | 98.7% | *Xyleborinus saxesenii* | 99.8 | *Xyleborinus saxesenii* |

## Discussion

sangerFlow derived taxa IDs matched with those derived from manual analysis. Our test data included Sanger reads with a wide range of sequencing quality, yet sangerFlow performed consistently producing exact same taxa IDs like those from the manual analysis. This suggested that sangerFlow was an automated, reproducible, and scalable bioinformatics pipeline for analysing Sanger PCR amplicon sequencing data. sangerFlow is the first of its kind as we are not aware of any command line-based bioinformatics pipeline that analyses Sanger-sequencing based PCR forward and reverse reads towards identifying the corresponding species. sangerFlow also has a broad application in the area of diagnosis and biosecurity surveillance as it can be applied to identify any pest and pathogen using a PCR plus Sanger sequencing assay (Fig. 3). Furthermore, sangerFlow removes the software installation requirements for individual analysis, which often involve software version conflicts and troubleshooting. sangerFlow achieves this by leveraging Nextflow pipeline manager and Singularity containers. In addition, hosting sangerFlow in GitHub along with a well-documented user manual allows the users to implement it with minimal computational expertise, a feature that is highly sought-after in the high-throughput diagnosis and surveillance laboratories.

**Figure 3.**
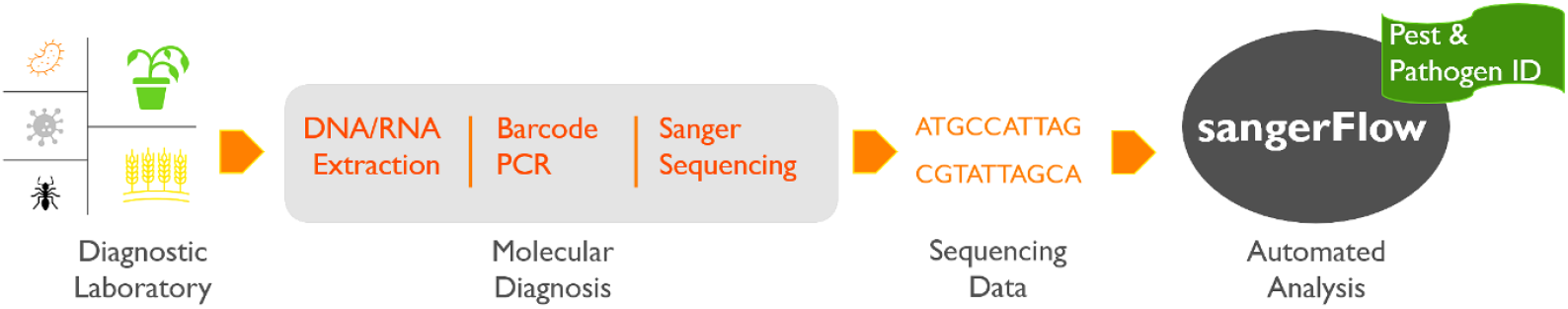
sangerFlow enables high-throughput automatic molecular diagnosis of pest and pathogens.

## Supplementary data

Supplementary data are available in the sangerFlow GitHub page (https://github.com/asadprodhan/sangerFlow).

## Funding

This work was supported by the DPIRD Diagnostics and Laboratory Services, Department of Primary Industries and Regional Development, Western Australia, Australia.

## Data availability statement

The data underlying this article are available in the sangerFlow GitHub page (https://github.com/asadprodhan/sangerFlow).

## Author contributions

M.A.P. conceptualised, developed, and validated the bioinformatics pipeline, sangerFlow. M.P. applied the pipeline to process large-scale in-house Sanger sequencing data including the test datasets. M.K. provided with feedback on the pipeline. M.A.P. wrote the manuscript. M.K. and M.P. commented on the manuscript. All authors read and approved the manuscript.

## Acknowledgements

We thank Dr Brenda Coutts and Mr Luc Shepherd (DPIRD Diagnostics and Laboratory Services, Department of Primary Industries and Regional Development) for the encouragement to develop a bioinformatics pipeline to tackle with the large influx of samples from the state-wide pest surveillance program. We thank the Pawsey Supercomputer Centre (https://pawsey.org.au/) for the computing resources.

## Declarations

### Ethics approval and consent to participate

Not applicable.

### Consent for publication

Not applicable.

### Humans or human data or any biological materials

Not applicable.

### Ethical approval

Not applicable.

### Competing interests

The authors declare that they have no competing interests.

### Availability and Requirements

Project name: sangerFlow

Project home page: https://github.com/asadprodhan/sangerFlow

Operating system: Linux

Programming language: Bash

Other requirements: Nextflow version 23.10.0, Singularity version 3.8.6, and Conda

License: GNU GPL v3 License

Any restrictions to use by non-academics: None

## Notes

### Competing Interest Statement

The authors have declared no competing interest.

